# Differential Drug Susceptibility Across Trichomonasvirus Species Allows for Generation of Varied Isogenic Clones of *Trichomonas vaginalis*

**DOI:** 10.1101/2024.08.12.607652

**Authors:** Carrie A. Hetzel, Akua A. Appah-Sampong, Austin R. Hurst-Manny, Max L. Nibert

## Abstract

*Trichomonas vaginalis* (Tvag) is a sexually transmitted human pathogen that is commonly infected with strains of one or more of five known species of Trichomonas vaginalis viruses (TVVs), members of genus *Trichomonasvirus*. TVVs are thought not to have an extracellular phase to their lifecycle and instead to be transmitted vertically from mother to daughter cells. As a result, generation of isogenic virus-positive and virus-negative sets of Tvag clones has been a major barrier to study interactions between TVVs and their host. Nucleoside analog 2’-C-methylcytidine (2CMC) has been recently reported to clear trichomonads of infections with TVV1, TVV2, and TVV3. We used 2CMC to treat a panel of Tvag isolates that collectively harbor at least one representative strain of each TVV species and thereby provided evidence that infections with TVV4 and TVV5 can also be cleared by 2CMC. Furthermore, our results suggest a newly identified difference in drug susceptibility between TVV species. We took advantage of these susceptibility difference to generate isogenic sets of Tvag clones harboring different combinations of the five TVV species. These results provide both new insight into differences between these species and new avenues for generating tools to study the potential roles of TVVs in Tvag biology.

## INTRODUCTION

Trichomonasviruses constitute a genus of double-stranded (ds)RNA viruses, genus *Trichomonasvirus*, comprising five currently recognized species: *Trichomonasvirus vagiprimus, Trichomonasvirus vagisecundus, Trichomonasvirus vagitertius, Trichomonasvirus vagiquartus*, and *Trichomonasvirus vagiquintus*. [1–4] These viruses are commonly referred to as Trichomonas vaginalis viruses (TVVs), and members of the five individual species are referred to as strains of TVV1, TVV2, TVV3, TVV4, and TVV5. They were previously classified in family *Totiviridae* but were recently moved to new family *Pseudototiviridae*, along with several other genera of dsRNA viruses that infect either fungi or protozoa [https://ictv.global/taxonomy]. Trichomonasviruses persistently infect the pathogenic protozoan *Trichomonas vaginalis* (abbreviated here as Tvag), an obligate human parasite and the causative agent of trichomoniasis, which tends to be more highly symptomatic in the female urogenital tract. [5] Trichomoniasis is the most common nonviral sexually transmitted infection (STI) worldwide.[5] In 2020, for example, there were estimated to be 156 million new cases per year in people aged 15-49.[6]

Trichomonasviruses are thought to increase the severity of trichomoniasis by inducing increased pelvic inflammation.[7] Clinically, infection by TVV-containing trichomonads is associated with higher degrees of inflammation and more severe disease progression.[8] Cervical epithelial cells show higher degrees of interferon-mediated inflammation when infected with TVV-containing parasites or purified TVV1 virions *ex vivo*. These cells can sense TVVs via toll-like receptor 3 (TLR3), which recognizes dsRNA. TLR3 triggers a proinflammatory cascade through interferon regulatory factor 3 (IRF3) and NFkB pathways.[7] The resulting increase in inflammation is thought to contribute to premature delivery during pregnancy, susceptibility to other sexually transmitted pathogens such as HIV and HPV, and other complications. [9] The inflammatory response may conceivably benefit the parasite by providing a microenvironment in the urogenital tract that promotes its growth, survival, and/or transmission.

Unlike well-known human viruses, TVVs are thought not to have an extracellular phase in their lifecycle and instead are transmitted endogenously from mother to daughter cells as the parasite divides.[10] Studying the effect of TVVs on the parasite host and the human super host has thus been historically difficult, as an uninfected Tvag isolate cannot simply be infected with virus by exogenous means in the laboratory. Previous studies have relied on comparisons between virus-positive and virus-negative Tvag isolates, but differences in the genetic backgrounds of these host trichomonads could well have confounded the results. Moreover, multiple TVV species can coinfect the same trichomonad, adding difficulty to the study of individual viruses.[10–12] Most studies to date have been performed using Tvag isolates that were singly infected with TVV1, singly infected with TVV3, or multiply infected with two or more TVV species. [7,13,14] To date, no isolate has been reported to be singly infected with either TVV4 or TVV5.

Generation of isogenic clones by clearing trichomonads of TVV infection has also proved challenging, as these viruses persist long term in the parasite, even after six months or more of serial passaging.[10] Furthermore, trichomonasviruses are thought to have few if any cytopathic effects on the protozoan. Combined, the various findings suggest to us that there is an evolutionary benefit for the protozoan to maintain the virus at tolerable levels. However, how this détente between virus and host is established and maintained, and what benefits these viruses might indeed provide to the host, have yet to be determined. Careful studies of isogenic clones of the parasite with and without the virus should help us to uncover such hypothesized benefits of the virus–host interactions.

A recent study by Narayanasamy et. al. [15] demonstrated that the nucleoside analogue 2’-C-methylcytidine (2CMC) is effective at reducing abundance of representative TVV1, TVV2, and TVV3 strains, and succeeded in using this strategy to generate cured clones of Tvag isolate T1c1, which had previously harbored TVV1. Here, we expand on the work done by those investigators and demonstrate that 2CMC is effective also against representative strains of TVV4 and TVV5. In addition, we elucidate species-specific differences in 2CMC susceptibility and use these differences to generate isogenic sets of singly infected, multiply infected, and uninfected clones of the same Tvag parent isolate.

## MATERIALS AND METHODS

### Tvag isolates and cell culture

This study used five Tvag isolates: JH37A#2, JH191A#4, JH32A#4, JH162A#4, and RU357. These isolates were purchased from the American Type Culture Collection (ATCC) and stored at -80 °C until use. Upon thawing, isolates were passaged every 1–3 days in Diamond’s Modified Media (DMM).[16] To ensure a clonal population, approximately 20 trichomonads were plated in 10 mL of 3% agar in DMM a 6-well culture plate. Individual trichomonads were then allowed to form colonies for 3-4 days and three colonies were transferred to liquid media upon becoming visible in soft agar. Resulting clones were labeled as A/B/C (e.g., JH37A#2-A) This process was then repeated to further ensure clonal purity. Resulting clones were designated A1/B1/A2/B2, etc.

### RNA extraction, cDNA synthesis, and qPCR

Total RNA was isolated using Trizol Reagent (Thermo Fisher Scientific) per manufacturer’s instructions. Resulting RNA was then incubated at 95 °C for 1 min before being placed immediately on ice to separate RNA strands. 200 µL of RNA was then used for cDNA synthesis using LunaScript RT SuperMix Kit (New England Biolabs). Resulting cDNA was next used as input for a quantitative (q)PCR reaction using Luna Universal qPCR Master Mix (New England Biolabs). Cycling conditions were as follows: 95 °C for 60 s, followed by 40 cycles of 95 °C for 15 s and 60 °C for 60 s. 96-well plates were run in an Applied Biosystems 7500 Fast Real Time PCR System (Invitrogen). Detection primers are listed in Table S1. Actin was used as a housekeeping control, and relative RNA expression was quantified using the 2-ΔΔCt method for normalizing gene expression.

### Drug treatment assays

In all assays, 5×10^5^ trichomonads were cultured in 10 mL DMM in a closed 15-mL conical tube. Cells were treated with 0–10 µL of 10 mM 2CMC dissolved in DMSO (final concentration 0–10 µM 2CMC) and supplemented with DMSO as a vehicle control to a final concentration of 0.1% DMSO. For time course experiments, cultures were passaged every 48 h by adding 1 mL of existing culture to 9 mL of fresh DMM and adding 2CMC and DMSO as needed to match starting concentrations. At each passage, 1 mL of culture was pelleted by centrifugation and resuspended in 1 mL of Trizol Reagent for RNA isolation, as described in section 2.2. For experiments involving varying 2CMC concentration, trichomonads were pelleted and resuspended in Trizol Reagent after 24 h.

### Generation of isogenic clones

After eight days of drug treatment, both treated and control cultures were plated in soft agar to generate clonal populations using the protocol described in section 2.1. Liquid cultures were then passaged approximately six times before RNA was extracted to determine if viral RNA was present as described in section 2.2. Clearance was defined a relative RNA expression of less than 0.1% of starting levels.

## RESULTS

### Treatment with 2CMC is effective at reducing RNA expression by strains of all five TVV species, but to varying degrees

In this study, we used Tvag isolates JH37A#2, JH191A#4, JH32A#4, JH162A#4, and RU357. Each isolate was clonally purified by picking colonies in soft agar twice serially to ensure a more homogeneous starting population of Tvag cells (clones designated A1, A2, B1, B2, etc.). RT-qPCR was then used to validate that spontaneous clearance of virus had not occurred. In doing so, we showed that JH37A#2-B1 and JH191A#4-B1 are singly infected with TVV1, JH32A#4-A1 is singly infected with TVV3, JH162A#4-A1 is multiply infected with TVV1, TVV2, TVV3, and TVV5, and RU357-B1 is multiply infected with TVV1, TVV2, TVV4, and TVV5 (Figure S1). Strains of the different TVV species present in each isolate are summarized in Table 1.

**Table 1.** TVV species present in Tvag isolates, verified by RT-qPCR (Figure S1).

| Tvag Isolate | TVV1 | TVV2 | TVV3 | TVV4 | TVV5 |
| --- | --- | --- | --- | --- | --- |
| JH37A#2-B1 | + | — | — | — | — |
| JH191A#4-B1 | + | — | — | — | — |
| JH32A#4-A1 | — | — | + | — | — |
| JH162A#4-A1 | + | + | + | — | + |
| RU357-B1 | + | + | — | + | + |

To date, 2CMC has been reported to inhibit strains of TVV1, TVV2, and TVV3, but has not been tested on strains of TVV4 and TVV5. [15] To corroborate and expand on these results, we treated Tvag isolates JH37A#2-B1, JH191A#4-B1, JH32A#4-A1, JH162A#4-A1 and RU357-A1 with 10 µM 2CMC or an equivalent amount of DMSO as a vehicle control. After eight days of passaging the trichomonads in media supplemented with 2CMC or DMSO, RNA was extracted, and viral RNA expression was measured by RT-qPCR. Across all of these Tvag isolates, RNA abundance from each TVV strain was reduced by greater than 99.8% relative to the respective control culture, including for TVV4 and TVV5 strains, which had not been shown before (Figure 1). Interestingly, reduction in viral RNA abundance seemed to differ between TVV species. In particular, the TVV1 and TVV2 strains in these Tvag isolates appeared to be less susceptible to 2CMC treatment at 48 h post-treatment than did the TVV3, TVV4, and TVV5 strains.

**Figure 1.**
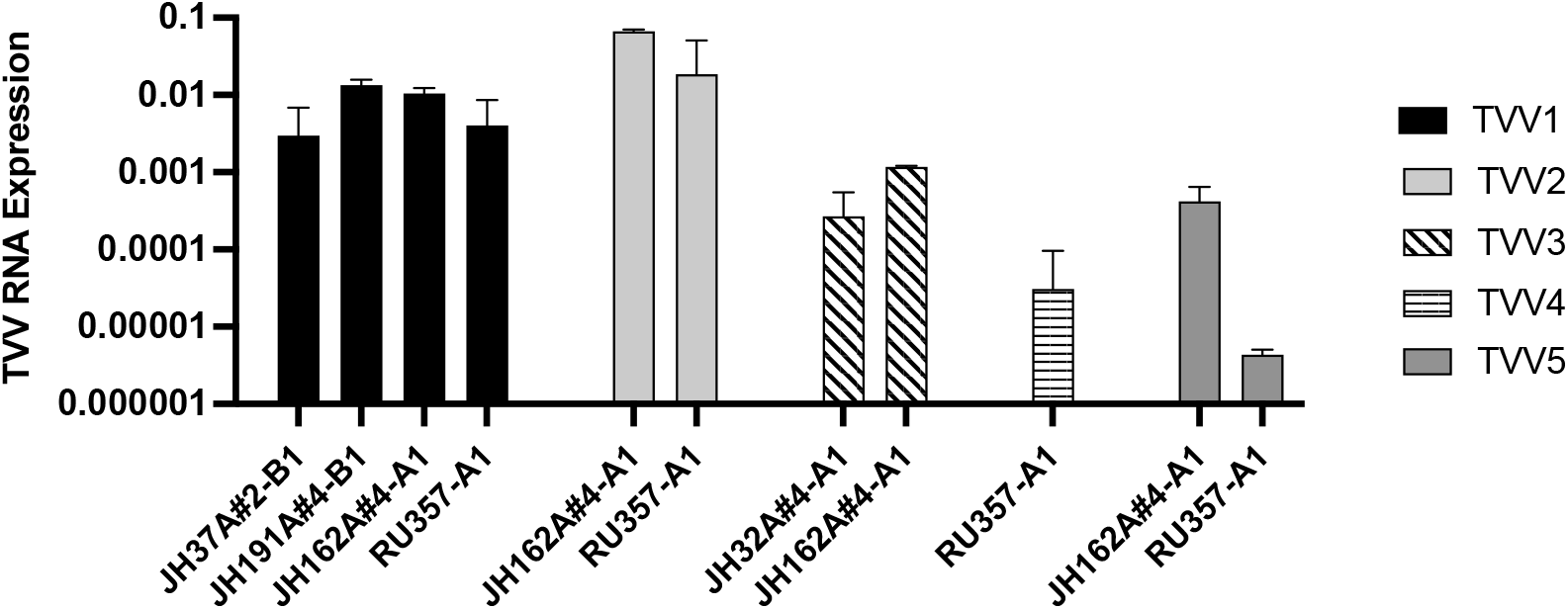
Treatment with 2CMC is effective at reducing RNA from strains of all five TVV species. Trichomonads were treated with 10 µM 2CMC for two days and relative viral RNA abundance was quantified using RT-qPCR. Expression values were calculated relative to a DMSO vehicle control and normalized to actin RNA expression as a housekeeping gene. Bars represent the mean of three technical replicates and error bars represent the standard deviation.

### Susceptibility to clearance by 2CMC varies by TVV species

To further investigate these apparent differences in 2CMC susceptibility, we passaged two of our singly infected Tvag isolates in 2CMC-containing media for eight days, monitoring viral RNA expression every two days (Figure 2A). In doing so, we saw that clearance of TVV3 RNA occurred much more quickly than clearance of TVV1 RNA. Moreover, TVV1 RNA was still detectable after eight days, while TVV3 RNA abundance had been reduced to below our limit of detection (cycle threshold (Ct) value > 35) by day 6. We also performed the same eight-day time course experiment using multiply infected Tvag isolate JH162A#4-A1 and saw that TVV3 RNA and TVV5 RNA reached undetectable levels by day 4, while TVV1 RNA and TVV2 RNA were still detectable at day 8, indicating a lower degree of susceptibility to inhibition by 2CMC on the part of TVV1 and TVV2 (Figure 2B). These findings are consistent with those in Figure 1 and were further corroborated by experiments with varying 2CMC concentrations (Figure 2C). The results in Figure 2 with multiply infected isolate JH162A#4-A1 are particularly interesting because monitoring reduction of TVV RNA abundance from different species in the same isolate eliminated the possibility of host differences confounding comparisons between isolates.

**Figure 2.**
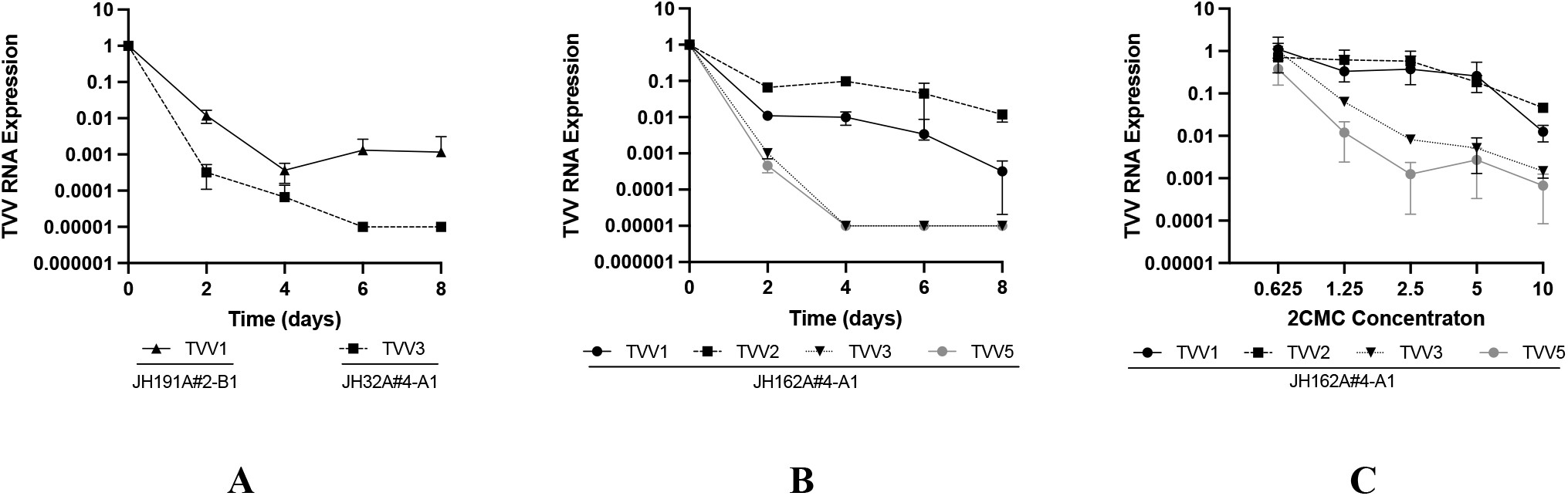
Susceptibility to 2CMC treatment varies between TVV species. Cells from (A) singly infected Tvag isolates JH191A#4-B1 and JH32A#4-A1 and (B) multiply infected isolate JH162A#4-A1 were passaged in media containing 10 µM 2CMC for eight days with timepoints every 48 h. (C) Cells from multiply infected isolate JH162A#4-A1were incubated in media containing 0–10 µM 2CMC for 24 h. Relative viral RNA abundance in each sample was quantified using RT-qPCR. Expression values were calculated relative to a DMSO vehicle control and normalized to actin RNA expression as a housekeeping gene. Datapoints represent the mean of three technical replicates and error bars represent the standard deviation.

To investigate if 2CMC susceptibility was consistent between strains of the same TVV species, we performed eight-day time course experiments with all four of the Tvag isolates that are infected with TVV1 (see Table 1). These four strains of TVV1 exhibited similar clearance rates, and TVV1 RNA was still detectable at day 8 for each of them (Figure 3A). We also compared the effect of varying 2CMC concentration on the two Tvag isolates that are singly infected with TVV1 (see Table 1) and saw little or no apparent difference in 2CMC susceptibility between these two TVV1 strains (Figure 3B), providing further evidence that 2CMC susceptibility is largely consistent across different strains of the same TVV species.

**Figure 3.**
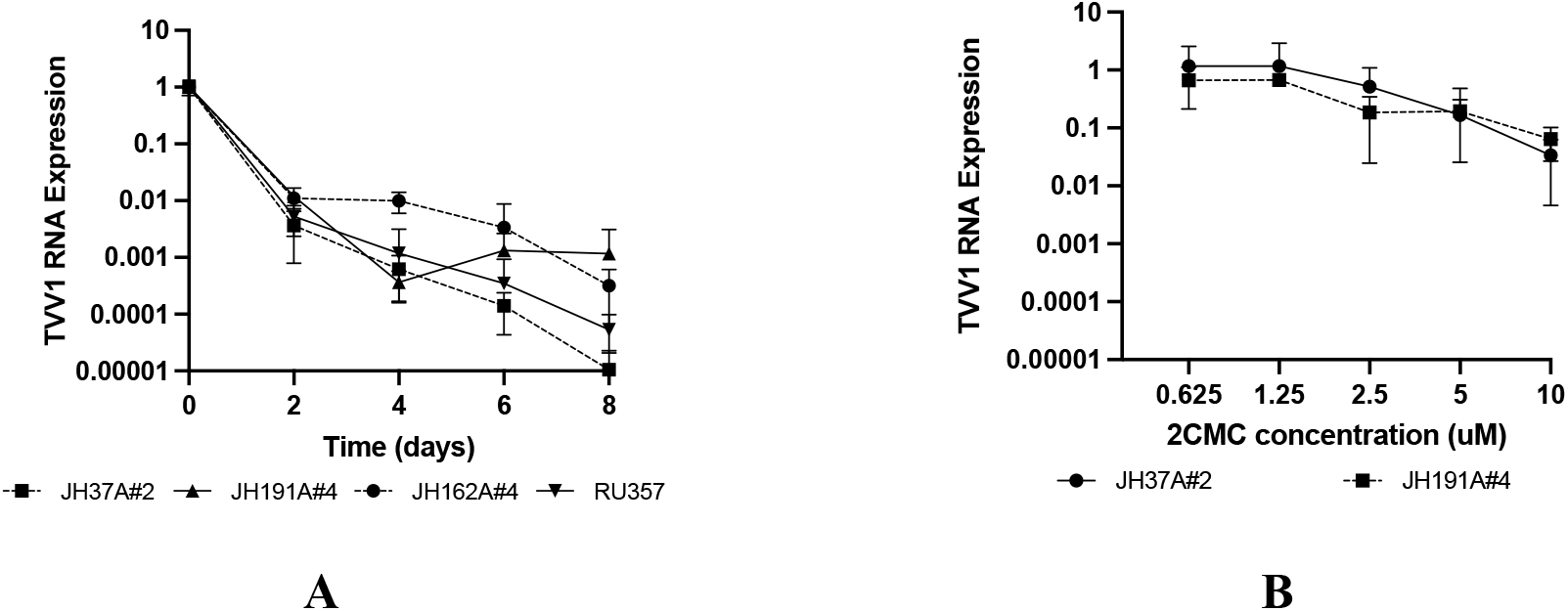
Susceptibility to 2CMC is similar between strains of the same TVV species. (A) Cells from the TVV1-infected isolates JH37A#2-B1, JH191A#4-B1, JH162A#4-A1, and RU357-B1 were passaged in media containing 10 µM 2CMC for eight days with timepoints every 48 h. (B) Cells from the singly TVV1-infected isolates JH37A#2-B1 and JH191A#4-B1 were incubated in media containing 0–10 µM 2CMC for 24 h. Relative viral RNA abundance in each sample was quantified and displayed as described for Figure 2.

Because TVV RNA was still present in some Tvag isolates at day 8 post-treatment, we anticipated that viral rebound would occur when 2CMC was removed at day 4. Furthermore, we were interested to see if the apparent differences in susceptibility between strains of different TVV species would affect the magnitude of rebound. When Tvag isolate JH37A#2-B1 (singly infected with TVV1) was tested in this manner, we saw that viral RNA rebounded to starting levels within two days after 2CMC removal (Figure 4A), despite viral RNA abundance having been reduced to ∼0.2% of starting levels at day 4. On the other hand, when this experiment was performed with Tvag isolate JH32A#4-A1 (singly infected with TVV3), we saw that viral RNA again rebounded within two days after 2CMC removal, but to only ∼7% of starting levels (Figure 4B), despite viral RNA abundance having been reduced in this case to ∼0.02% of starting levels (nearly the limit of detection) at day 4. Thus, consistent with the greater magnitude of TVV3 clearance from Tvag isolate JH32A#4-A1 at day 4 of 2CMC treatment, TVV3 RNA abundance did not rebound to the same degree in that isolate as did TVV1 RNA in isolate JH37A#2-B1. Sanger sequencing of the coding regions of both virus populations after rebound revealed no mutations, suggesting that rebound was not the result of outgrowth of resistant mutants, but rather the outgrowth of the wild-type viruses that had persisted in the face of 2CMC treatment for four days.

**Figure 4.**
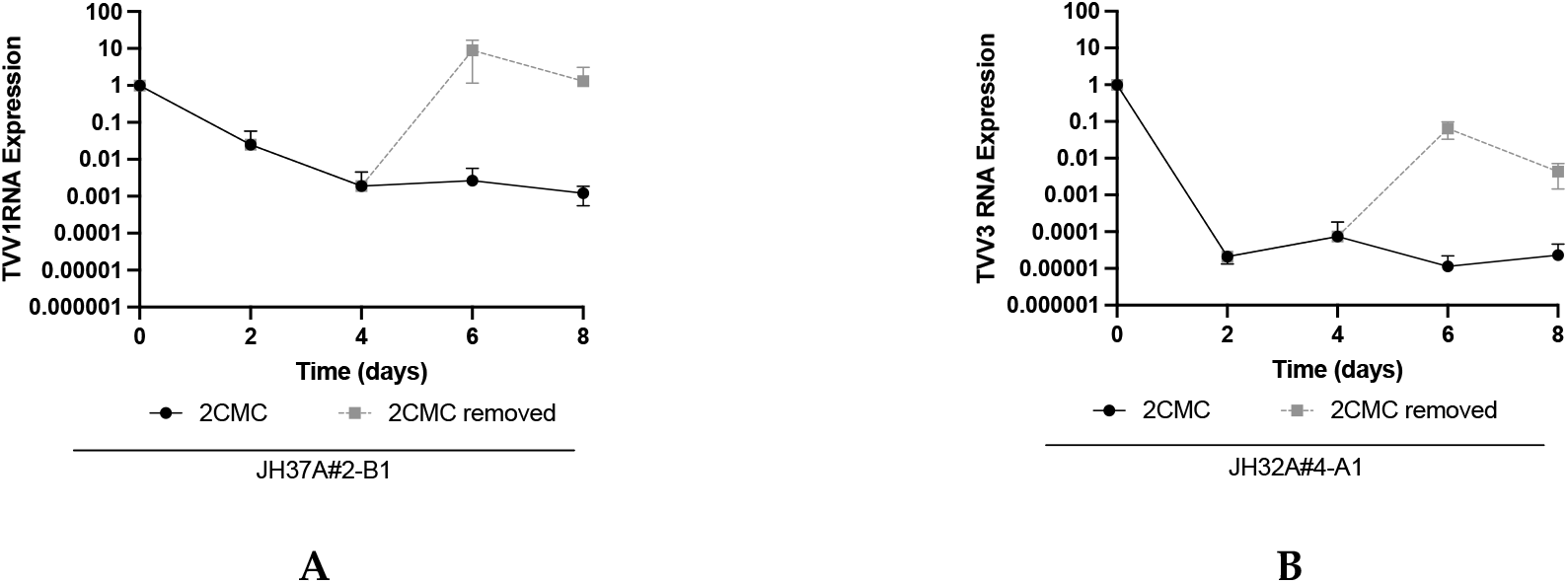
TVV infection persists after four days of 10 µM 2CMC treatment and rebounds upon 2CMC removal. Trichomonads from (**A**) singly TVV1-infected Tvag isolate JH37A#2-B1 and (**B**) singly TVV3-infected Tvag isolate JH32A#4-A1 were passaged in media containing 10 µM 2CMC for four days. At day 4 for each isolate, 2CMC was removed from one series of samples and continued for the other. Relative RNA abundance at each timepoint was quantified and displayed as described for Figure 2.

### Isogenic clones generated after 8 days of 2CMC treatment

We wanted to generate isogenic uncured and cured clones of Tvag isolates, as was done by Narayanasamy et al. [15], to use for future experiments. However, due to our results regarding time to clearance and viral rebound, we passaged our isolates in 2CMC or DMSO, as a vehicle control, for eight days before plating for Tvag colonies in soft agar to obtain clonal populations. To ensure these clones were cured of TVV, we passaged each approximately six times in culture after selection and monitored viral RNA expression. In doing so, we were able to generate isogenic sets from singly infected isolates JH37A#2-B1, JH191A#4-B1, and JH32A#4-A1 (Figure 5A-C). For these experiments, we defined a cured strain as having viral RNA abundance reduced to 0.1% or less of starting levels, i.e., reduced by three orders of magnitude or more, following six or more passages in culture. All of the clones we derived from 2CMC treatment of isolates JH191A#4-B1 and JH32A#4-A1 were thereby considered cured. Interestingly, one of the four 2CMC-treated clones from JH37A#2-B1 (clone 7 in Figure 5A) remained infected with TVV1, providing further evidence that viral RNA can persist in at least some Tvag cells in a culture even after eight days of 2CMC treatment. Sanger sequencing of this clone revealed four synonymous mutations in the coding region of TVV1, but these mutations did not appear to affect 2CMC susceptibility relative to the original virus (Figure S2), suggesting to us that these mutations were not responsible for the persistence of TVV1 in this clone.

**Figure 5.**
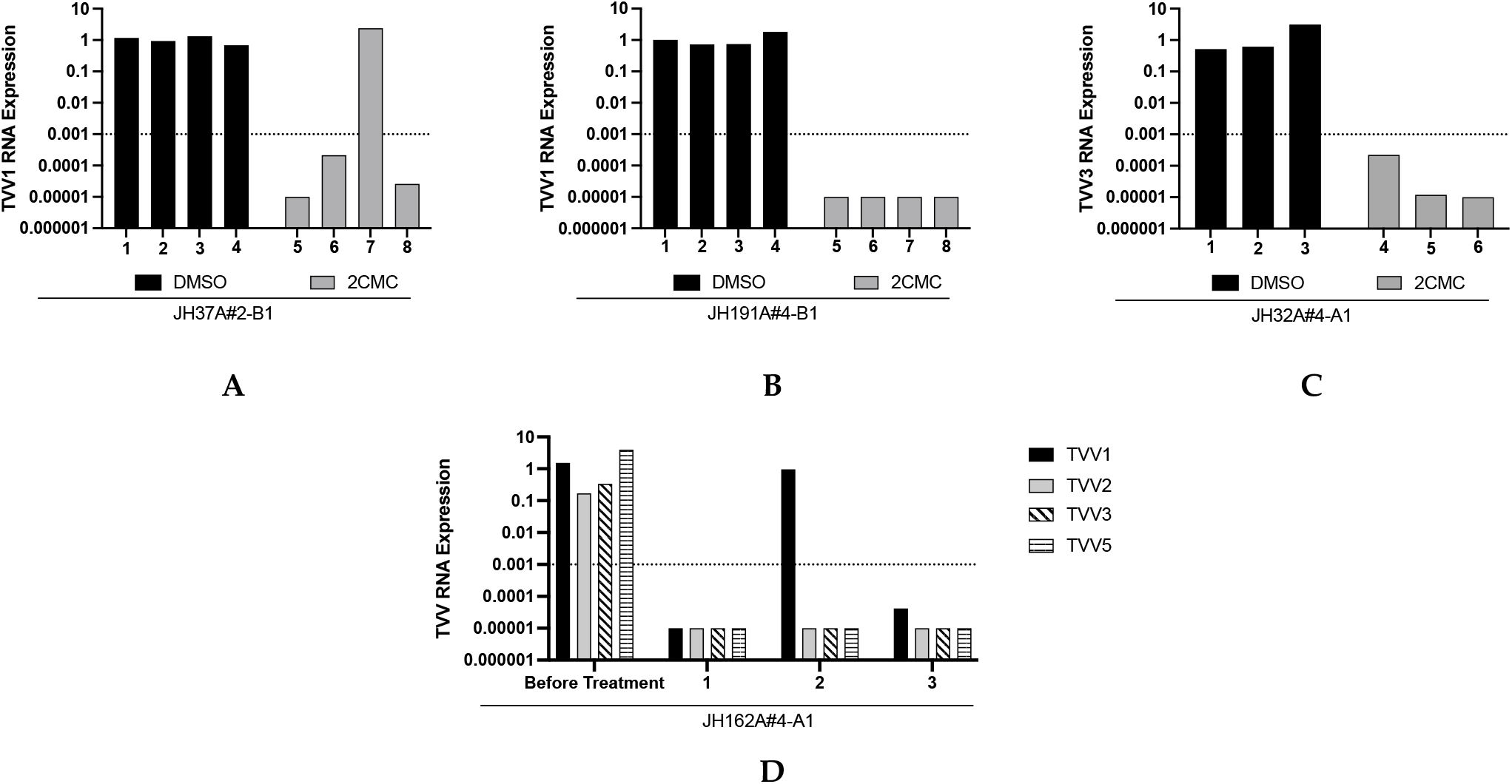
Treatment with 10 µM 2CMC allows for clearance of virus from singly and multiply infected isolates. Trichomonads from isolates (**A**) JH37A#2-B1, (**B**) JH191A#4-B1, (**C**) JH32A#4-A1, and (**D**) JH162A#4 were passaged in media containing 10 µM 2CMC or the equivalent amount of DMSO for eight days. At day 8, trichomonads were plated in soft agar and allowed to form colonies. Three or four individual colonies for each isolate and treatment condition were screened for the presence of trichomonasviruses by RT-qPCR. Relative viral RNA abundance in each sample was quantified as described for Figure 1. Dotted line represents viral RNA abundance below which clones were considered to be cured. In **D**, the samples labeled “Before Treatment” reflect analysis of the parent Tvag isolate (not DMSO treated and cloned in parallel to the 2CMC treatment yielding clones 1–3).

In addition to our singly infected Tvag isolates, we treated multiply infected isolate JH162A#4-A1 in the same manner as above and screened three clones after eight days of 2CMC treatment. Of these three clones, one remained singly infected with TVV1, while the other two were completely cured of TVV infections (Figure 5D). The presence of TVV1 in clone 2 is consistent with our results in Figures 1 and 2, which provided evidence that TVV1 strains are less susceptible to 2CMC inhibition than are TVV3 and TVV5 strains. Sanger sequencing revealed no mutations in the coding regions of the TVV1 strain in clone 2, suggesting to us that the persistence of TVV1 in this clone did not reflect the selection of a 2CMC-resistant mutant. Interestingly, we did not detect TVV2 in any of the three clones, despite the lesser susceptibility of TVV2 strains to 2CMC inhibition shown in Figure 1 and Figure 2. However, selecting for singly TVV2-infected or TVV1/TVV2-coinfected clones may be possible with further screening.

## 4. Discussion

Trichomonasviruses constitute an understudied genus, despite their potential as a model system to investigate virus–host interactions in the context of long-term viral persistence in a human pathogen, *T. vaginalis*. This is in part due to the challenge of working with such viruses that appear to lack an extracellular phase in their lifecycle. The generation of virus-positive and virus-negative Tvag clones derived from the same parent isolate, to use for comparative analyses, is thus crucial. Narayanasamy et al. [15] demonstrated the capacity of 2CMC to clear representative strains of TVV1, TVV2, and TVV3 and used this approach to generate uncured and cured isogenic Tvag clones. In this study, we extended those previous results to demonstrate that 2CMC is also effective in clearing representative strains of TVV4 and TVV5 from Tvag isolates, allowing for generation of a wider range of isogenic pairs of Tvag clones. Furthermore, we demonstrated the possibility of using 2CMC treatment to create isogenic sets of Tvag clones containing varied combinations of strains of the five TVV species and generated the first reported isogenic set containing singly infected, multiply infected, and uninfected Tvag clones derived from the same parent isolate (see Figure 5). To date, little is known about the relative impact of single vs. multiple TVV infections on the protozoan host, and so such isogenic sets of Tvag clones are essential for allowing careful investigations of these differences.

Isogenic sets of clonally purified host cell cultures are additionally useful for related studies of virus–host biology in the numerous other systems involving protozoan, fungal, or other hosts that are persistently infected by viruses that are not routinely transmitted by extracellular means, i.e., including but extending beyond trichomonasviruses and *T. vaginalis*. Trichomonasviruses are an example of viruses that provide an intriguing model for studying the virus–host arms race in the context of long-term viral persistence. It is largely unknown how the détente between endosymbiotic viruses and their hosts is achieved and maintained, and how that détente can be broken. Moreover, because Tvag is an early-diverging eukaryote [17], understanding this détente might provide valuable information about the evolution of host defenses, as Tvag lacks many classical innate immune pathways [18] that might be expected to participate in this détente. Isogenic sets of Tvag clones harboring different combinations of trichomonasviruses may, for example, allow for differential gene expression analyses to help to uncover host biochemical pathways that might be involved in controlling TVV replication.

Our results additionally demonstrated the importance of using extended treatment of Tvag cultures with 2CMC in order to achieve viral clearance from all or nearly all Tvag cells in the cultures, followed by clonal purification of cured cells. When we removed 2CMC after four days of treatment, strains of both TVV1 and TVV3 rebounded, despite having been reduced to nearly undetectable levels in the case of TVV3 (see Figure 4). Sanger sequencing revealed that the coding regions of the TVV strains present in the rebounded cultures were identical to the starting sequences, indicating to us that rebound was primarily explained by outgrowth of wild-type viruses that had persisted in many of the Tvag cells after four days of 2CMC treatment, and not by outgrowth of any 2CMC-resistant mutants that may have been selected. Based on the relative levels of rebound, we interpreted that after four days of treatment, TVV1 had persisted in most Tvag cells in isolate JH37A#2-B1, whereas TVV3 had persisted in < 10% of Tvag cells in isolate JH32A#4-A1. Similarly, strains of TVV1 that remained after eight days of 2CMC treatment in one clone each from Tvag isolates JH37A#2-B1 and JH162A#4-A1 (see Figure 5A and D) provided further evidence for the persistence of virus in at least some Tvag cells after extended treatments with 2CMC.

Currently, there are no reported Tvag isolates that are singly infected with either TVV4 or TVV5. Further investigations might be fruitful for identifying inhibitors to which strains of particular TVV species are uniquely resistant, which would allow for the generation of Tvag clones that are singly infected with strains of each of the different species, including TVV4 and TVV5. Nucleoside analogs in general are well-known inhibitors of viral polymerases [19], and thus further study of differential susceptibility of TVV strains to a larger panel of nucleoside analogs would seem to be especially promising. Because TVV1 and TVV2 strains exhibited the least susceptibility to 2CMC inhibition, they currently present a limitation to being cleared from Tvag cells of a multiply infected parent isolate before strains of the other species in that isolate have been cleared. Thus, inhibitors that are specifically more effective against TVV1 and/or TVV2 strains are especially needed to allow an even greater number of combinations of isogenic clones to be generated.

In addition to further study into the relationships between virus (TVV) and host (Tvag), as well as superhost (human), future directions of this work will be to elucidate why the apparent differences in 2CMC susceptibility between TVV species exist. According to analyses performed by Manny et. al. [20], sequences from TVV2 and TVV5 strains are more similar to one another than to strains of the other species, as also are sequences from TVV3 and TVV4 strains, with TVV1 strains being the most divergent from the others. Intriguingly, susceptibility to 2CMC treatment did not appear to replicate this pattern in our experiments. Instead, TVV1 and TVV2 strains appeared to be less susceptible to inhibition by 2CMC than did TVV3, TVV4, and TVV5 strains. No obvious patterns of sequence differences have been noted in our analyses to date to try to explain this result. We have also performed preliminary structure predictions using AlphaFold2 [21] for the viral polymerase from strains of the different TVV species, but no obvious structural differences to explain these differences in 2CMC susceptibility have yet been recognized. Of further note, apparent susceptibility to 2CMC by strains of different TVV species did not appear to correlate with starting levels of viral RNA in the Tvag parent cultures. For example, in Tvag isolate JH162A#4-A1, TVV3 appeared to have the highest relative starting level of RNA abundance, followed by TVV1, then TVV5, then TVV2 (see Figure S1).

To conclude, further work is needed in a number of important areas identified by this study as suggested above. In addition, structural determinations of the viral polymerase from the different TVV species could allow greater insight into how nucleoside analog 2CMC may interact with this protein, which is essential for viral replication. Lastly, identification of novel strains or clones of the different TVV species, with increased or decreased susceptibility to 2CMC achieved either by further sampling of naturally occurring variation among Tvag clinical isolates or by selecting 2CMC-resistant mutants from Tvag isolates in the laboratory, could also help to elucidate how 2CMC may interact with the viral polymerase and provide additional information about the differences between TVV species. It is even conceivable, though perhaps unlikely, that the effect of 2CMC occurs via a less direct mechanism, such as by affecting nucleotide substrate levels in Tvag cells, which then differentially impact the five TVV species.

