## Supplementary Tables and Figures for "Differential Drug Susceptibility Across Trichomonasvirus Species Allows for Generation of Varied Isogenic Clones of *Trichomonas vaginalis*"

**Table S1. Primers used for RT-qPCR**

| Target | Forward Primer Sequence | Reverse Primer Sequence |
|---|---|---|
| Actin | TCACAGCTCTTGCTCCACCA | AAGCACTTGCGGTGAACGAT |
| TVV1-JH37A#2 | ATTAGCGGCGTTTGTGATGCA | CCTGGGGTTTGCGTTCCTTG |
| TVV1-JH191A#4 | ATTAGCGGTGTTTGTGATGCA | TTGCCATGCTCTAGCTTGCG |
| TVV3-JH32A#4 | GAAGCTGAGCTTCTCGTCACAG | ATGAGGTTGGACAGACTTCCTGTC |
| TVV1-JH162A#4 | ATTAGCGGTGTTTGTGATGCA | TTGCCATGCTCTAGCTTGCG |
| TVV2-JH162A#4 | CTGACTTACACCGACAGTTGGAC | GTCTTTTAAGAAAGCATCGTTGCGAC |
| TVV3-JH162A#4 | GATTGGTGCATCGCTAGCATTG | TTGGTTGCCACTCCCATGATG |
| TVV5-JH162A#4 | TCGTCTCTGTCTAGCTGCCTCT | CGTTCTTGCACCAGAATGGTGATG |
| TVV1-RU357 | ATTAGCGGCGTTTGTGATGCA | ACTTGAGGCTTGCATTCCTTGAG |
| TVV2-RU357 | CTGACTTACACCGACAGTTGGAC | GTCTTTTAAGAAAGCATCGTTGCGAC |
| TVV4-RU357 | GCCGACTTGAAGGTCAACTGC | GTGTAGATAGTTCTTATGGCGAGACGC |
| TVV5-RU357 | CCTATATGCTCGTCTCTGTCTGGC | GAATGGACGTGGTCAGTGAAACTG |

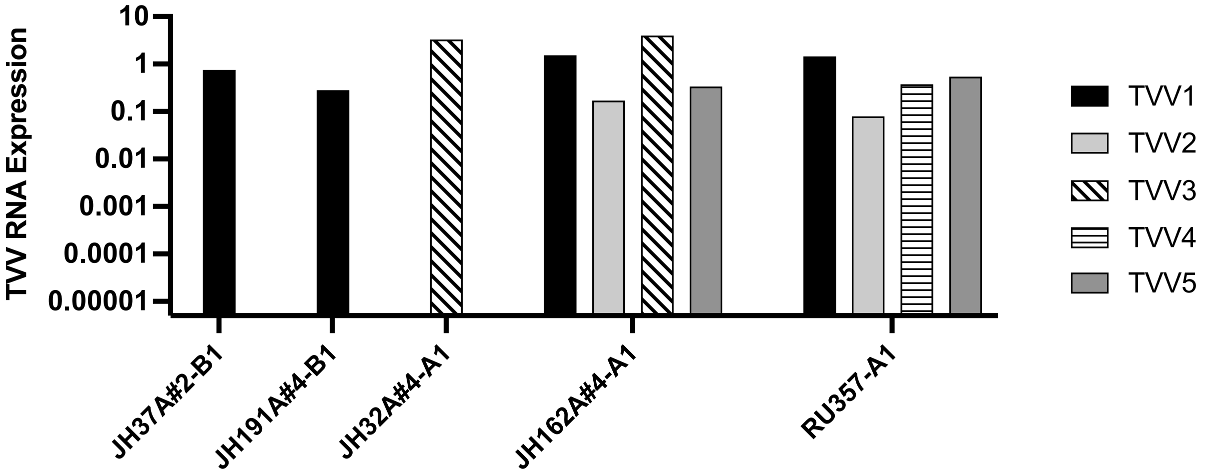


**Figure S1. Validation of the presence of trichomonasviruses in isolates tested.** RNA was extracted from each culture and screened for the presence of virus by RT-qPCR as described in section 2.1.

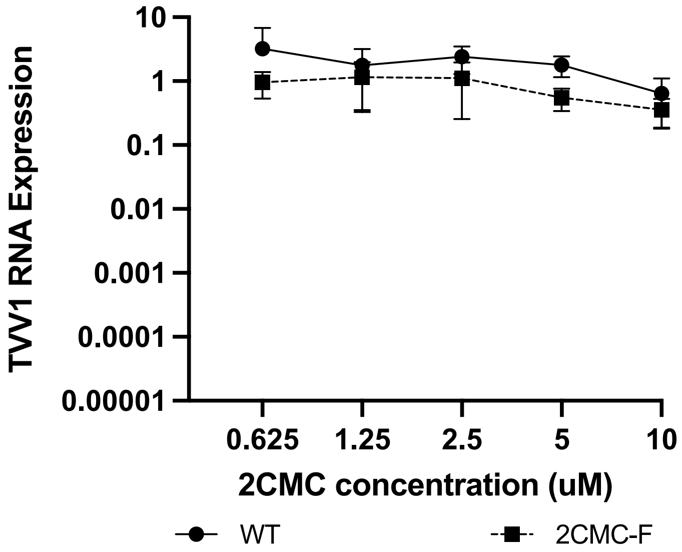


**Figure S2. Susceptibility to 2CMC is not significantly different between TVV1 in parent isolate JH37A#2-B1 and TVV1 in 2CMC-treated but uncured clone JH37A#2-B1-2.** Cells from parent isolate JH37A#2-B1 and clone JH37A#2-B1-2 were incubated in media containing 0–10 µM 2CMC for 24 h. Relative viral RNA abundance in each sample was quantified and displayed as described for Figure 2.
